# Efficient Alcoholic Conversion of Glycerol by Engineered *Saccharomyces cerevisiae*

**DOI:** 10.1101/2021.05.24.445540

**Authors:** Sadat M. R. Khattab, Takashi Watanabe

## Abstract

Glycerol is an eco-friendly solvent that enhances plant biomass decomposition via glycerolysis in many pretreatment methods. Nonetheless, the lack of efficient conversion of glycerol by natural *Saccharomyces cerevisiae* hinders its use in these methods. Here, we have aimed to develop a complete strategy for the generation of efficient glycerol-converting yeast by modifying the oxidation of cytosolic nicotinamide adenine dinucleotide (NADH) by an O_2_-dependent dynamic shuttle, while abolishing both glycerol phosphorylation and biosynthesis. By following a vigorous glycerol oxidation pathway, the engineered strain increased the conversion efficiency (CE) to up to 0.49 g ethanol/g glycerol (98% of theoretical CE), with production rate > 1 g·L·h, when glycerol was supplemented in a single fed-batch fermentation in a rich medium. Furthermore, the engineered strain fermented a mixture of glycerol and glucose, producing > 86 g/L bioethanol with 92.8% CE. To our knowledge, this is the highest ever reported titer in this field. Notably, this strategy changed conventional yeast from a non-grower on minimal medium containing glycerol to a fermenting strain with productivity of 0.25−0.5 g·L·h and 84−78% CE, which converted 90% of the substrate to products. Our findings may improve the utilization of glycerol in several eco-friendly biorefinery approaches.

**IMPORTANCE:** With the development of efficient lignocellulosic biorefineries, glycerol has attracted attention as an eco-friendly biomass-derived solvent that can enhance the dissociation of lignin and cell wall polysaccharides during the pretreatment process. Co-conversion of glycerol with the sugars released from biomass after glycerolysis increases the resources for ethanol production and lowers the burden of component separation. However, titer productivity hinders the industrial application of this process. Therefore, generation of efficient glycerol-fermenting yeast will promote the applicability of integrated biorefineries. Furthermore, glycerol is an important carbon source for the production of various chemicals. Hence, control of the metabolic flux of yeast grown on glycerol will contribute to the generation of cell factories that produce chemicals such as ethanol from glycerol, which will boost biodiesel and bioethanol industries. Additionally, the use of glycerol-fermenting yeast will reduce global warming and generation of agricultural waste, leading to the establishment of a sustainable society.

---

The demand for ethanol has increased remarkably because of its use in sanitizers in the COVID-19 pandemic, necessitating enhancement in its production via various pathways, including glycerol fermentation. In the past decade, glycerol (C_3_H_8_O_3_)-producing industries, particularly the biodiesel industry, have produced substantial amounts of glycerol (1). However, glycerol is reduced more than other fermentable sugars (2). In addition, glycerol as a carbon source is poorly utilized, primarily via the glycerol-3-phosphate pathway (hereafter referred to as the G3P pathway), which is composed of glycerol kinase (*GUT1*) and flavin adenine dinucleotide (FAD)-dependent mitochondrial glycerol-3-phosphate dehydrogenase (*GUT2*) (3). Yeast prefers biosynthesizing glycerol, as it mitigates osmotic stress and optimizes redox balance (4). In contrast, glycerol catabolism is subject to repression and transcriptional regulation of glucose metabolism via several respiratory factors, as well as *GUT1* and *GUT2* (5-7).

Several groups have attempted to ferment glycerol by overexpressing the native oxidative pathway (DHA; dihydroxyacetone pathway) in *S. cerevisiae*. Overexpression of the endogenous glycerol dehydrogenase (*ScGCY1)* with dihydroxyacetone kinase (*ScDAK1*) led to the production of 0.12 g ethanol/g glycerol (g^e^/g^g^), and the production rate was 0.025 g·L·h (8). The importance of glycerol as a carbon source that can be utilized by yeast cells prompted a study on 52 *S. cerevisiae* strains, which revealed the intraspecies diversity of yeast, ranging from good glycerol growers to non-growers. The growth phenotype is controlled by quantitative traits (9). Reports showed that many genes such as *UBR2*, encoding a cytoplasmic ubiquitin-protein ligase E3, link *GUT1* with growth on glycerol in synthetic medium in the absence of supplements (10). In contrast, heterologous replacement of G3P with DHA, combined with the use of the glycerol facilitator *FPS1*, resulted in the restoration of growth characteristics similar to those of the parental strain or even higher (11). This replacement in a glycerol grower ancestor by limiting the oxygen in shaking flask cultures increased ethanol production from 0.165 to 0.344 g^e^/g^g^ in buffered synthetic medium. The maximum titer produced reached 15.7 g/L after 144 h (12). Until now, glycerol has been reported as a non-fermentable carbon source for *Saccharomyces cerevisiae*, and attempts have been made to ferment it using this yeast (13). Furthermore, the methylotrophic yeast *Ogataea polymorpha* has been modified for bioethanol production from glycerol by overexpressing multiple genes involved in either the DHA or G3P pathways. In addition, *FPS1*, pyruvate decarboxylase (*PDC*) 1, and alcohol dehydrogenase (*ADH*) 1 were overexpressed. Nonetheless, the overall titer produced was 10.7 g/L ethanol with a conversion efficiency 0.132 g^e^/g^g^ (14). However, reports on known native or genetically engineered *S. cerevisiae* strains that ferment or convert glycerol efficiently to ethanol are lacking.

We have previously reported microwave-assisted pretreatments of recalcitrant lignocellulosic biomass in aqueous glycerol (15), which was further improved with the catalysis of alum [AlK(SO_4_)_2_] (16). Fractionation of sugars from glycerol is expensive. Therefore, *S. cerevisiae* capable of efficiently fermenting glycerol with glucose after glycerolysis pretreatment of biomass should be developed. Therefore, in this study, we have aimed to model the *S. cerevisiae* to redirect glycerol traffic to bioethanol production in the presence of glucose via systematic metabolic engineering outlined in Fig. 1. Modeling the conversion of glycerol and sugars after lignocellulosic biomass glycerolysis will synergize ethanol production, which is necessary for cost-effective distillation processes with fewer purification steps. Thence, many biorefineries are promoted, as well as the merger of the bioethanol and biodiesel industries. [“ (Figure 1)”]

**Fig. 1.**
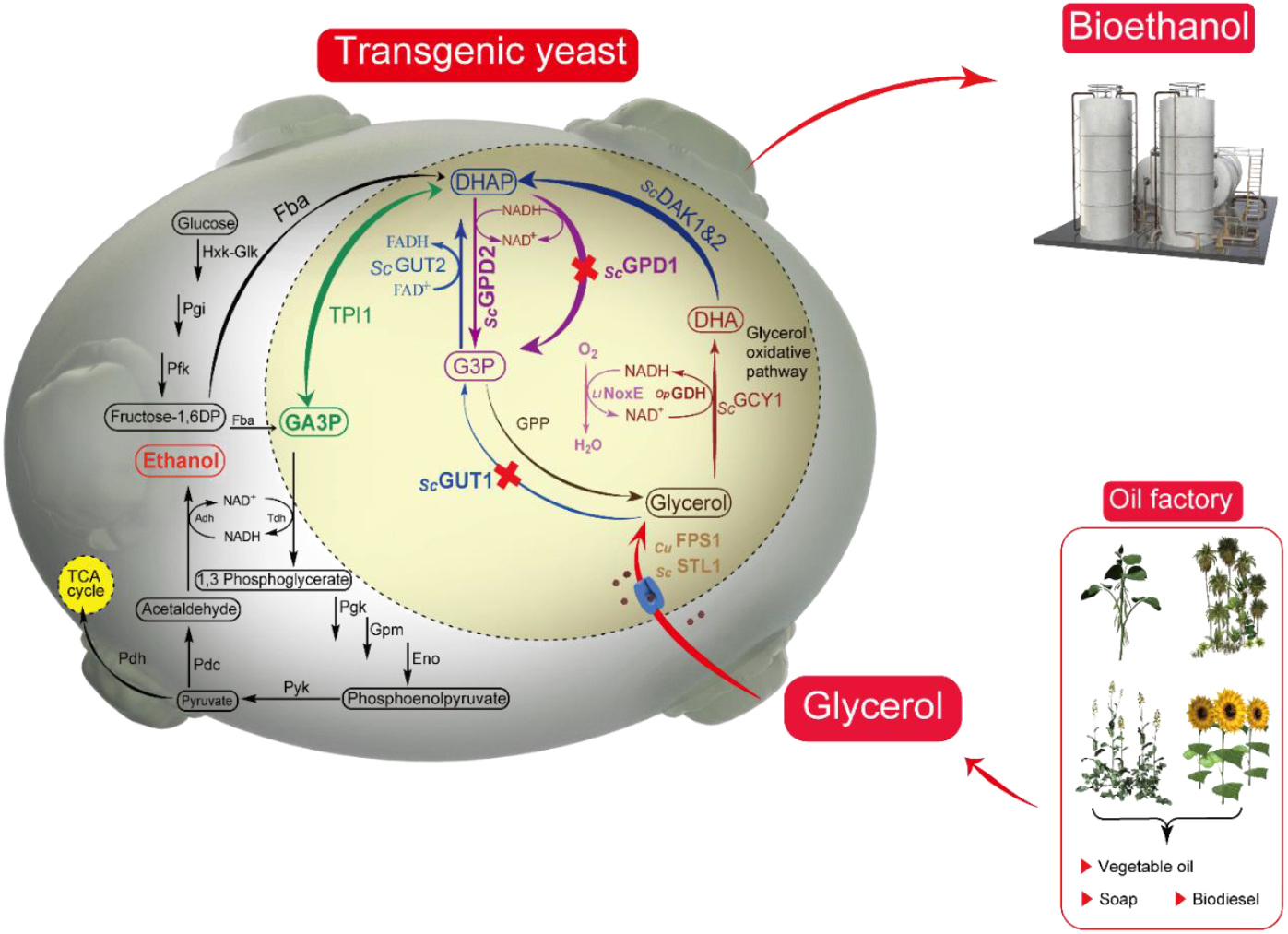
Schematic showing the integrative scenario of a biorefinery with new generation of glycerol-fermenting yeast and redirection of glycerol influxes to ethanol production in *Saccharomyces cerevisiae* via retrofitted native glycerol anabolic and catabolic pathways using the robust oxidative route with restoration of the NAD^+^ cofactor via O_2_-dependent dynamics of water-forming NADH oxidase. During pathway re-routing, glycerol-3-phosphate dehydrogenase 1 (*ScGPD1*) and glycerol kinase 1 (*GUT1*) were knocked out. Highlighted circle indicates the overexpressed indigenous *S. cerevisiae* enzymes dihydroxyacetone kinase (*ScDAK*) 1 and 2, as well as triosephosphate isomerase (*ScTPI*) 1, heterologous glycerol dehydrogenase from *Ogataea polymorpha* (*OpGDH*), glycerol facilitator from *Candida utilis* (*CuFPS1*), and water-forming NADH oxidase from *Lactococcus lactis* subsp. *lactis* Il1403 (*LlNoxE*).

## RESULTS AND DISCUSSION

### Growth of the original strain on glycerol as the sole carbon source

*S. cerevisiae* cannot ferment glycerol. In addition, some strains of *S. cerevisiae* cannot grow in culture media containing glycerol as the sole carbon source in the absence of supplements. Swinnen et al. classified the S288C strain as a negative glycerol grower on synthetic medium without supporting supplements (9). *S. cerevisiae* D452-2 (Table 1), which was used in this study, is an ancestor of S288C (17-19) (Fig. S1). The strain could not use glycerol as the sole carbon source in yeast nitrogen base (YNB) medium supplemented with 20 mg/L leucine and histidine, and 5 mg/L uracil or even when 10× supplementation was used. The maximum OD_600_ after 120 h was only 0.28 (Fig. S2). Hence, in this study, we used a conventional yeast peptone (YP) medium to avoid these limitations.

**Table 1.**
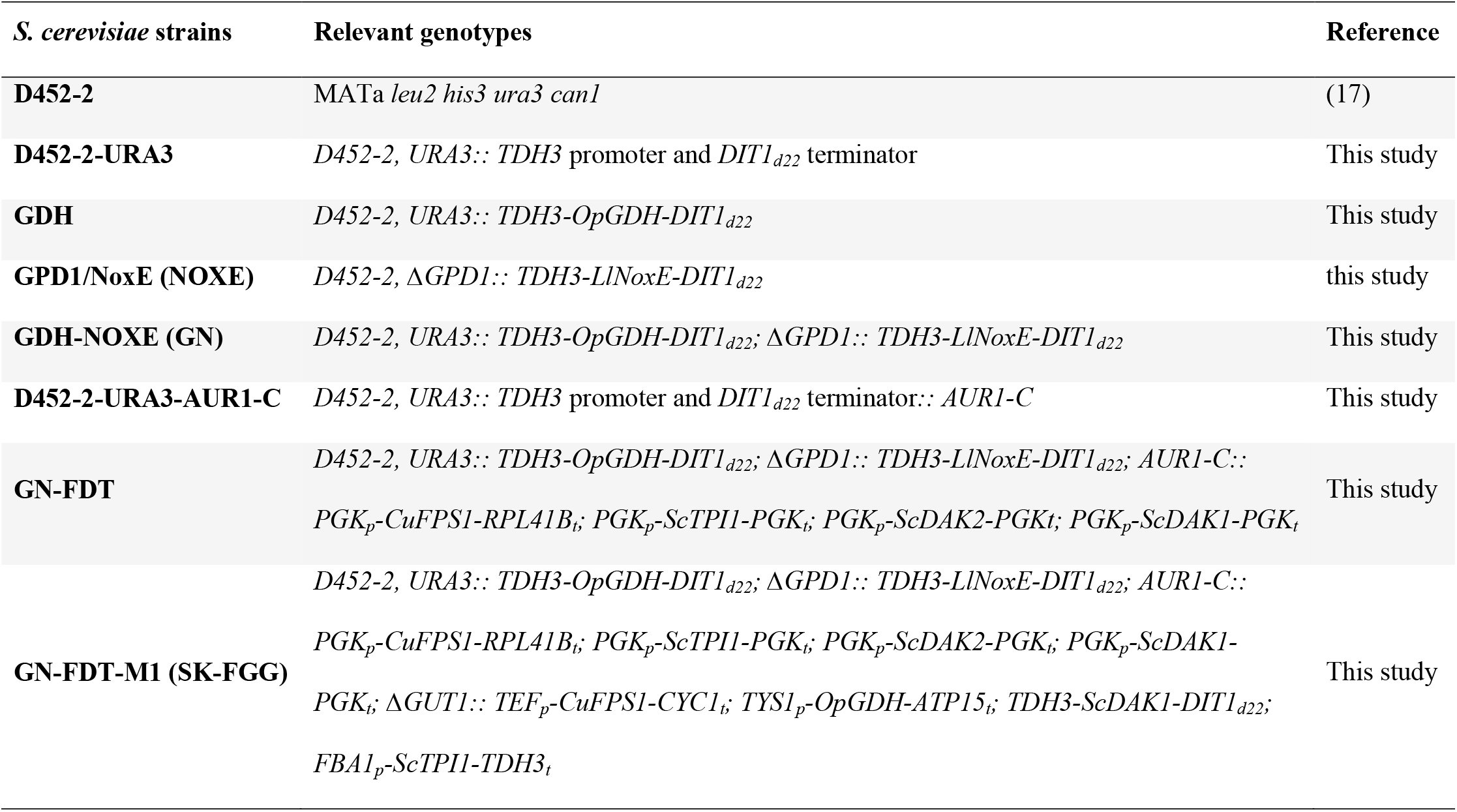
Characteristics of *Saccharomyces cerevisiae* strains generated in this study.

| *S. cerevisiae* strains | Relevant genotypes | Reference |
| --- | --- | --- |
| **D452-2** | MATa *leu2 his3 ura3 can1* | (17) |
| **D452-2-URA3** | D452-2, *URA3:: TDH3* promoter and *DIT1*<sub>d22</sub> terminator | This study |
| **GDH** | D452-2, *URA3:: TDH3-OpGDH-DIT1*<sub>d22</sub> | This study |
| **GPD1/NoxE (NOXE)** | D452-2, Δ*GPD1:: TDH3-LlNoxE-DIT1*<sub>d22</sub> | this study |
| **GDH-NOXE (GN)** | D452-2, *URA3:: TDH3-OpGDH-DIT1*<sub>d22</sub>; Δ*GPD1:: TDH3-LlNoxE-DIT1*<sub>d22</sub> | This study |
| **D452-2-URA3-AUR1-C** | D452-2, *URA3:: TDH3* promoter and *DIT1*<sub>d22</sub> terminator::*AUR1-C* | This study |
| **GN-FDT** | D452-2, *URA3:: TDH3-OpGDH-DIT1*<sub>d22</sub>; Δ*GPD1:: TDH3-LlNoxE-DIT1*<sub>d22</sub>; *AUR1-C:: PGK<sub>p</sub>-CuFPS1-RPL41B<sub>t</sub>; PGK<sub>p</sub>-ScTPII-PGK<sub>t</sub>; PGK<sub>p</sub>-ScDAK2-PGK<sub>t</sub>; PGK<sub>p</sub>-ScDAK1-PGK<sub>t</sub>* | This study |
| **GN-FDT-M1 (SK-FGG)** | D452-2, *URA3:: TDH3-OpGDH-DIT1*<sub>d22</sub>; Δ*GPD1:: TDH3-LlNoxE-DIT1*<sub>d22</sub>; *AUR1-C:: PGK<sub>p</sub>-CuFPS1-RPL41B<sub>t</sub>; PGK<sub>p</sub>-ScTPII-PGK<sub>t</sub>; PGK<sub>p</sub>-ScDAK2-PGK<sub>t</sub>; PGK<sub>p</sub>-ScDAK1-PGK<sub>t</sub>; Δ*GUT1:: TEF<sub>p</sub>-CuFPS1-CYC1<sub>t</sub>; TYS1<sub>p</sub>-OpGDH-ATP15<sub>t</sub>; TDH3-ScDAK1-DIT1*<sub>d22</sub>; *FBA1<sub>p</sub>-ScTPII-TDH3<sub>t</sub>** | This study |

### Enhancement of glycerol flux via glycerol dehydrogenase

To overcome the limitations of glycerol flux in *S. cerevisiae, OpGDH* from the methylotrophic yeast *O. polymorpha*, encoding glycerol dehydrogenase (20), was heterologously overexpressed in the *URA3* locus of D452-2 under the control of an efficient expression system (21) consisting of the glyceraldehyde 3-phosphate dehydrogenase promoter (*TDH3*) and mutated DITyrosine terminator (*DIT1*_*d22*_). In contrast to the native strain, which did not show any enzyme activity, 3.5 mU/mg GDH activity was observed for the transformant harboring *OpGDH* (Table 2). Compared to 167 mM potassium phosphate (pH 7.5), HEPES buffer increased glycerol dehydrogenase activity by 3.75-fold, which is similar to that reported previously (22). Consequently, compared to that of the reference strain, glycerol consumption increased from 8% to 58%. Concomitantly, the production of ethanol increased by 2.64-fold, from 4.47 g/L (in the reference strain) to 11.82 g/L, reaching productivity of 0.27g^e^/g^s^ after 26 h (Fig. 2). Aßkamp et al. reported that external mitochondrial NADH dehydrogenase (Nde1) is involved in glycerol metabolism in the presence of oxygen as the final electron acceptor (23), although cytosolic NAD^+^ was partially regenerated. Under semi-aerobic conditions, the NADH/NAD^+^ ratio increased by 45% (Table 2). Therefore, overexpression of *OpGDH* in *S. cerevisiae* is the first step in the alcoholic fermentation of glycerol. Nonetheless, overexpression of only *OpGDH* in GDH strain was not sufficient to induce an efficient conversion (Fig. 2). In contrast, the conversion efficiency in strains overexpressing endogenous glycerol dehydrogenase *ScGCY1* alone or in combination with whole DHA pathway was lower than that of strains overexpressing *OpGDH* (data not shown). [“ (Table 1)”]

**Table 2.**
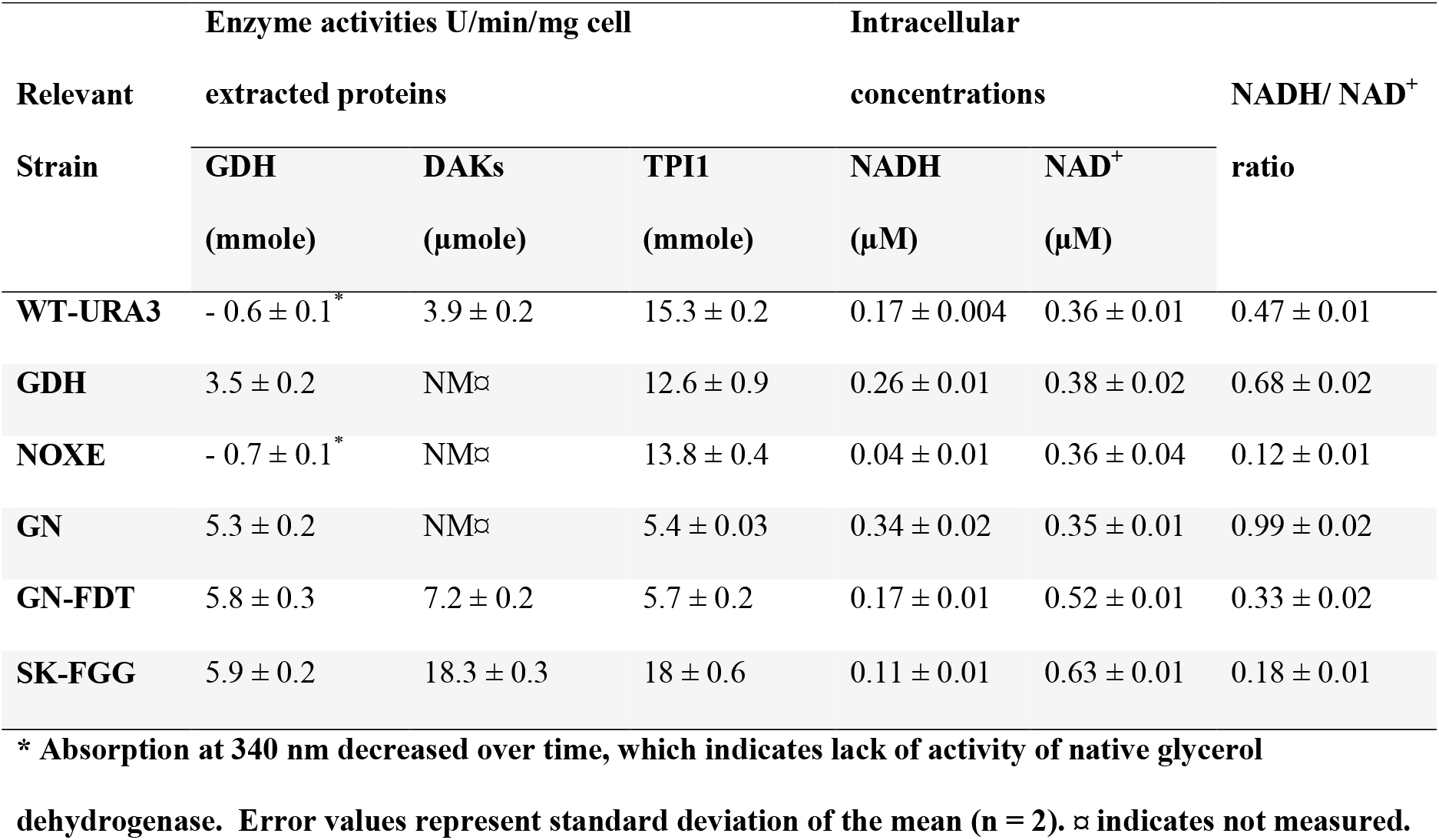
Specific activities of glycerol dehydrogenase (*GDH*), dihydroxyacetone kinase (*DAK*), and triosephosphate isomerase (*TPI1*), and NADH/NAD^+^ ratio with their intracellular concentrations in the recombinant strains used in this study.

| Relevant<br>Strain | Enzyme activities U/min/mg cell<br>extracted proteins |  |  | Intracellular<br>concentrations |  | NADH/ NAD<sup>+</sup><br>ratio |
| --- | --- | --- | --- | --- | --- | --- |
|  | GDH | DAKs | TPII | NADH | NAD<sup>+</sup> |  |
|  | (mmole) | (μmole) | (mmole) | (μM) | (μM) |  |
| **WT-URA3** | - 0.6 ± 0.1<sup>*</sup> | 3.9 ± 0.2 | 15.3 ± 0.2 | 0.17 ± 0.004 | 0.36 ± 0.01 | 0.47 ± 0.01 |
| **GDH** | 3.5 ± 0.2 | NM<sup>⌘</sup> | 12.6 ± 0.9 | 0.26 ± 0.01 | 0.38 ± 0.02 | 0.68 ± 0.02 |
| **NOXE** | - 0.7 ± 0.1<sup>*</sup> | NM<sup>⌘</sup> | 13.8 ± 0.4 | 0.04 ± 0.01 | 0.36 ± 0.04 | 0.12 ± 0.01 |
| **GN** | 5.3 ± 0.2 | NM<sup>⌘</sup> | 5.4 ± 0.03 | 0.34 ± 0.02 | 0.35 ± 0.01 | 0.99 ± 0.02 |
| **GN-FDT** | 5.8 ± 0.3 | 7.2 ± 0.2 | 5.7 ± 0.2 | 0.17 ± 0.01 | 0.52 ± 0.01 | 0.33 ± 0.02 |
| **SK-FGG** | 5.9 ± 0.2 | 18.3 ± 0.3 | 18 ± 0.6 | 0.11 ± 0.01 | 0.63 ± 0.01 | 0.18 ± 0.01 |

**\* Absorption at 340 nm decreased over time, which indicates lack of activity of native glycerol**
**dehydrogenase. Error values represent standard deviation of the mean (n = 2). ⌘ indicates not measured.**

**Fig. 2.**
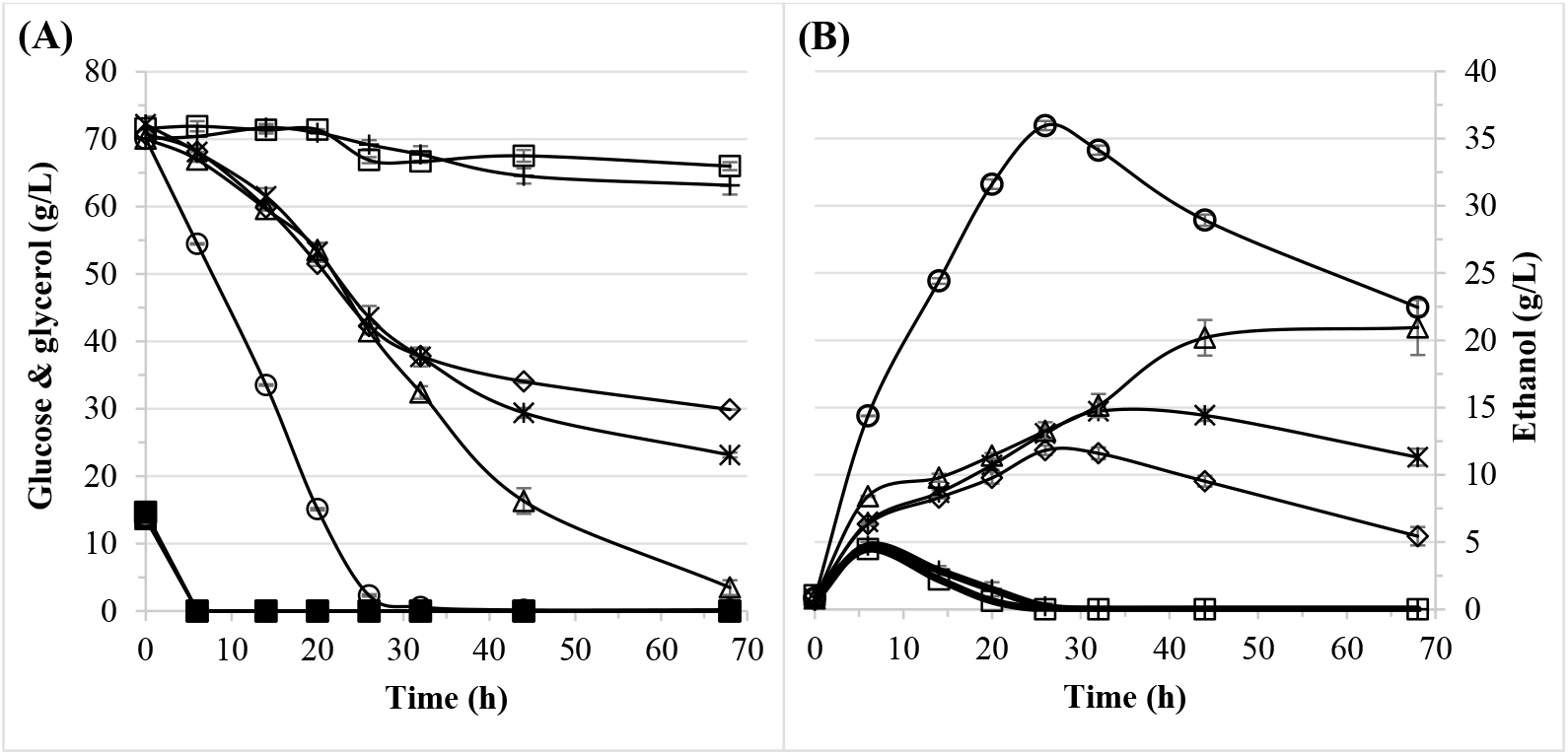
Comparison of the time course of glycerol-glucose fermentation between the *S. cerevisiae* strains used in this study. Reference ancestral strain (square); NOXE strain (plus); GDH strain (rhomboid); GN strain (asterisk); GN-FDT strain (triangle); GN-FDT-M1[SK-FGG] (circle). (a) Glucose consumption (closed symbols), glycerol consumption (opened symbols), and (b) ethanol production (opened symbols). Fermentation was performed under semi-aerobic conditions in flasks with 20:200 liquid medium: flask volume at 30 °C with shaking at 180 rpm. YP medium was supplemented with (w/v) 1.5% glucose and 6% glycerol. Data represent the average of three independent experiments that were running at the same time. Error bars represent the standard deviation of the mean.

### Modification of NADH oxidation

NADH is produced during glucose metabolism, and heterologous expression of *NoxE* in *S. cerevisiae* has been used to successfully decrease excess NADH production during fermentation of glucose, with increase in ethanol, 2,3-butanediol or acetoin production (24-28). We substituted glycerol-3-phosphate dehydrogenase (GPD) shuttles with *LlNoxE* and observed that this replacement abolished glycerol biosynthesis by 98%, with increase in ethanol production by 9% during fermentation of 10% glucose (data not shown). Replacement of *ScGPD1* by *LlNoxE* in the GDH strain overcame the deficiency of NADH oxidation by the natural cytosolic shuttles and induced efficient regeneration of NAD^+^ for glycerol oxidation. In addition, *ScGPD1* replacement reduced circulation of DHAP into glycerol biosynthesis or/and G3P—glycerolipid pathway. Subsequently, this promoted continuous conversion of glycerol to ethanol. Substantial improvement in ethanol production was observed using this combination in the GN strain (Table 1), which harbouring *OpGDH* and *LlNoxE*, with production being 28% of that of the GDH strain. The ethanol titer increased from 11.82 to 13.27 g/L (0.31g^e^/g substrate consumed) after 26 h, and then to 14.42 g/L with further extension of the fermentation time to 32 h (Fig. 2b). Surprisingly, the enzyme activity of glycerol dehydrogenase increased by 51%, whereas that of *TPI1* decreased by 65%. The reasons underlying these changes remain unclear. The effectiveness of oxidation was further evident from the restoration of the NADH/NAD^+^ ratio (Table 2). NAD^+^ level was restored during glycerol conversion by integrating *LlNoxE* in the GDH strain; nevertheless, it was not sufficient for efficient ethanolic conversion. [“ (Table 2)”]

### Complete overexpression of the DHA pathway

Considering the efficiency of glycerol conversion to ethanol by the GN strain, the activities of other genes in the DHA pathway, namely, *ScTPI1, ScDAK*s, and *CuFPS1*, may still be limited, which may affect the traffic of glycerol to ethanol. The DHA pathway is unusual, as DHAP is distributed in phospholipid and methylglyoxal biosynthesis pathways (29, 30). Therefore, further reduction of the bifurcation of DHAP can be overcome by overexpressing *ScTPI1*. There has been no report of *ScTPI1* overexpression aiming to enhance ethanolic fermentation, although the pivotal role of *ScTPI1* is evident from another study on glycerol production (31). Furthermore, dihydroxyacetone may accumulate inside cells expressing *OpGDH* and reduce transport of DHA, and its toxific effects (32) could be the driving force for alcohol production. *ScDAK1* and *CuFPS1* support the fermentation of glycerol to ethanol (8, 12, 14). In this study, we further supported the activities of *ScTPI1* and *ScDAK2* (with considerably lower *Km*_(DHA-ATP)_) with *ScDAK1* and *CuFPS1*. After integrating this set of genes in the GN strain, the GN-FDT strain unequivocally overcame the obstacle of ethanol re-consumption before the complete utilization of glycerol. The consumption rate reached 1 g·L·h, with increase in ethanol titer to 21 g/L, which represents 145% and 469% of the titers of the GN and native strains, respectively. Nonetheless, the conversion efficiency was less than 48% of the theoretical value (Fig. 2). Compared to that of the native reference strain, dihydroxyacetone kinase activity in GN-FDT improved by 185%, while compared to that of the GN strain, glycerol dehydrogenase and *ScTPI1* were activated by 9% and 6%, respectively (Table 2). Hence, increasing the activities of *ScDAK* and *ScTPI1* by supporting the influx of glycerol by *CuFPS1* represents another essential step for improving conversion of glycerol to ethanol. “ (Figure 2)”]

### Reinforcement of the DHA pathway while abolishing the native G3P route

Native *S. cerevisiae* adapts to redox regulation via glycerol biosynthesis (4). Furthermore, the activity of *ADH* within the cells grown on glycerol-yeast extract medium was ten times higher than that in cells cultured in glucose (33). In addition, the availability of ATP for *ScDAKs* may be limited by the presence of *ScGUT1* (11). Therefore, enhancement of the efficiency of glycerol conversion was expected by another copy of the DHA pathway concomitant with the replacement of *ScGUT1*. Accordingly, overexpressing another copy of *CuFBS1, OpGDH, ScDAK1*, and *ScTPI1* through the M1 module (Fig. 3), which were placed under a highly constitutive expression system (21, 34, 35), enhanced the activities of *ScDAK* and *ScTPI1* by 256% and 316%, respectively, compared to that in the GN-FDT strain. In addition, the efficiency of NADH oxidation increased with reduction in the NADH/NAD^+^ ratio to 0.11 (Table 2). Interestingly, the GN-FDT-M1 (SK-FGG) strain showed outstanding conversions; the glycerol consumption rate reached 2.6 g·L·h, and ethanol productivity was 1.38 g·L·h with conversion efficiency of 0.44 g^e^/g^s^ (Fig. 2). This suggests that the glycerol flux and efficiency of conversion are controlled by the extent to which the DHA pathway restores NAD^+^. “ (Figure 3)”]

**Fig. 3.**
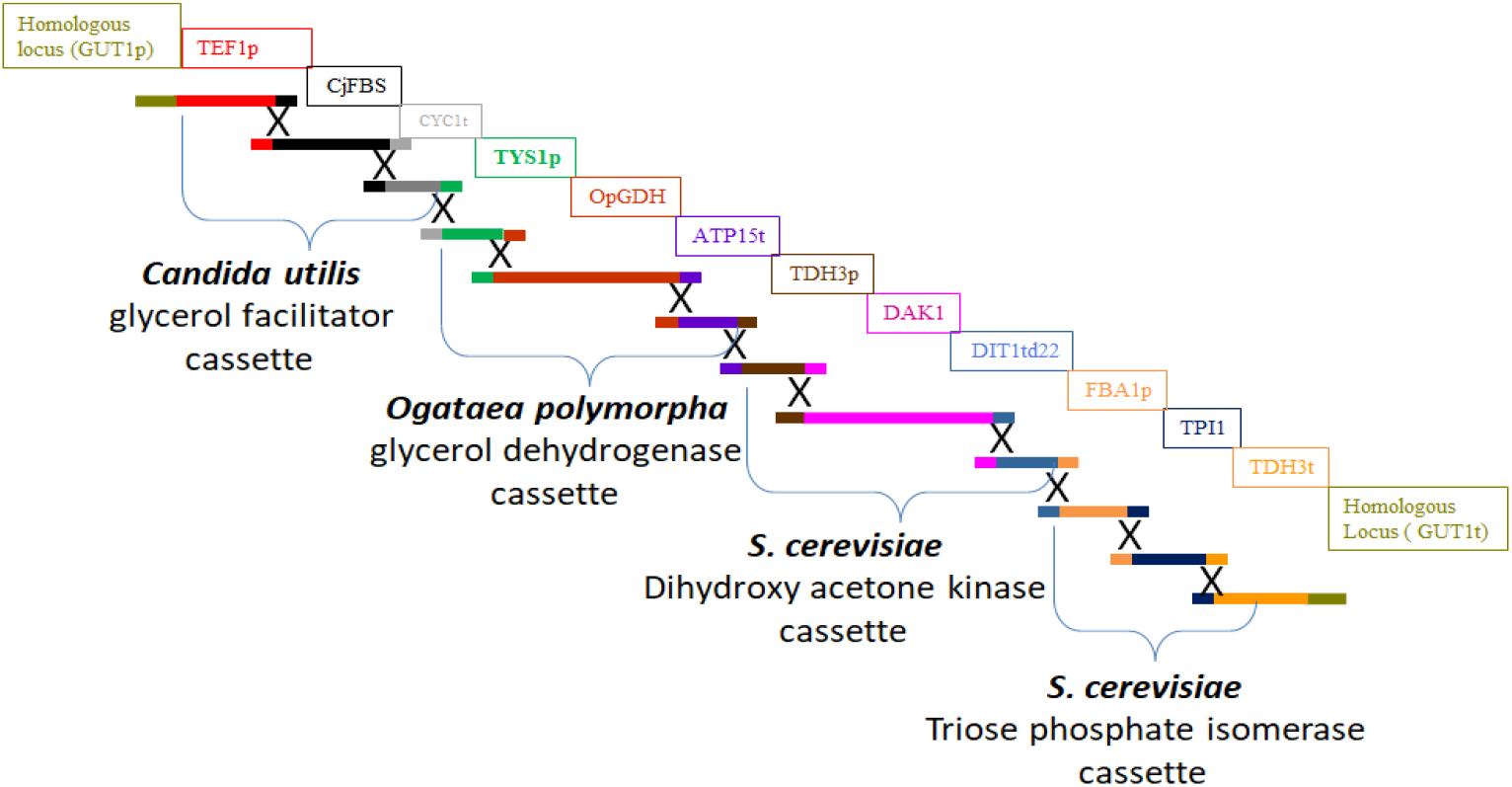
Module M1 for replacing *ScGUT1* via homologous recombination using multiplex CRISPR Cas 9.

To further investigate the metabolic engineering in SK-FGG, alcoholic conversion of higher initial concentrations (110 g/L) of glycerol in the presence or absence of 22.5 g/L glucose (Table 3 [A and B]) was tested. Furthermore, fed-batch fermentation of approximately 100 g/L glycerol was performed under condition [B] (Table 3 [C]). To further confirm that the engineered strain did not substantially repressed by glucose metabolism (5-7), the glucose level was doubled to 45 g/L with reduction in glycerol concentration by 25% to 82 g/L before the fed-batch fermentation of glycerol (Table 3 [D]). This scheme was also assessed for the ability to yield an economical distillable ethanol titer (Table 3 [D]). The state [D] is similar to that obtained after the glycerolysis process (16). With case [A], the conversion efficiency reached 98% (0.49 g^e^/g^g^) with production rate of >1 g·L·h of ethanol after consumption of 82.5 g/L glycerol under the fed-batch condition. In addition, 1.14 g/L acetic acid was produced. Even in the presence of 22.55 g/L glucose, the conversion efficiency of glycerol was almost unaltered (Table 3 [B]). Interestingly, the engineered strain was exceptional in its capacity of harmonizing the conversion of glycerol with glucose, along with an accumulation of > 86 g/L bioethanol during an additional fed-batch of glycerol (Table 3 [D]). Compared to that in case [C], cell density increased by 31%, along with a minor reduction in the efficiency of ethanol conversion in [D] (Table 3). [“ (Table 3)”]

**Table 3.** Fermentation characteristics of strain SK-FGG at high concentrations of glycerol in the absence and presence of glucose in fed-batch culture.

| Parameter | Fermentation conditions and products values |  |  |  |
| --- | --- | --- | --- | --- |
|  | One fed-batch | One fed-batch | Two fed-batch | Two fed-batch |
| **Fermentation time (h)** | 40 ± 0.30 | 40 ± 0.30 | 96 ± 0.30 | 96 ± 0.30 |
| **Total consumed glycerol (g/L)** | 82.5 ± 1.63 | 75.77 ± 1.4 | 141.88 ± 2.7 | 141.7 ± 5 |
| **Initial glycerol concentration (g/L)** | 111 ± 1 | 108.15 ± 1 | 108.17 ± 1 | 82.84 ± 1 |
| **Glycerol fed-batch concentration (g/L)** | NA | NA | 94 ± 1 | 99 ± 2.5 |
| **Total consumed glucose (g/L)** | NA | 22.5 ± 1.2 | 22.2 ± 0.67 | 44.7 ± 1 |
| **Rate of consumption (g·L·h)** | 2.1 ± 0.2 | 2.4 ± 0.1 | 1.7 ± 0.1 | 1.9 ± 0.1 |
| **Ethanol yield (g/L)** | 40.4 ± 0.1 | 48.2 ± 1.2 | 79.7 ± 1.1 | 86.5 ± 2.5 |
| **Rate of ethanol production (g·L·h)** | 1.01 ± 0.01 | 1.2 ± 0.02 | 0.83 ± 0.01 | 0.9 ± 0.02 |
| **Efficiency of ethanol production (g<sup>e</sup>/ g<sup>s</sup>)** | 0.49 ± 0.01 | 0.49 ± 0.01 | 0.48 ± 0.01 | 0.46 ± 0.01 |
| **Efficient/ theoretical (%) <sup>+</sup>** | 98 | 98 | 97.2 | 92.8 |
| **Acetic acid accumulation (g/L)** | 1.14 ± 0.02 | 1.12 ± 0.02 | 1.67 ± 0.1 | 2.46 ± 0.02 |
| **Total conversion (g<sup>p</sup>/ g<sup>s</sup>) <sup>++</sup>** | 0.5 ± 0.01 | 0.5 ± 0.01 | 0.5 ± 0.01 | 0.48 ± 0.01 |
| **Final cell density (OD<sub>600</sub>)** | 10.6 ± 0.2 | 11.6 ± 0.2 | 11.6 ± 0.3 | 15.2 ± 0.3 |

Oxygen availability in liquid medium: flask volume at 30 °C with shaking at 200 rpm (20:200).
(g<sup>e</sup>/g<sup>s</sup>) \*, gram ethanol per gram of glycerol. (g<sup>p</sup>/ g<sup>s</sup>) <sup>++</sup>; gram products per gram of substrate.
Efficiency (theoretical lrb%) <sup>+</sup>: C<sub>6</sub>H<sub>12</sub>O<sub>6</sub> → 2 C<sub>2</sub>H<sub>5</sub>OH + 2 CO<sub>2</sub> → **Eq. 1** C<sub>3</sub>H<sub>8</sub>O<sub>3</sub> + ½ O<sub>2</sub> → C<sub>2</sub>H<sub>5</sub>OH + CO<sub>2</sub> + H<sub>2</sub>O → **Eq. 2**
(1 g) → 0.51 g + 0.49 g (1 g) → 0.5 g + 0.478 g

### Ethanolic conversion of glycerol as the sole carbon source

To avoid the interference of nutrients in the complex YP medium, the conversion efficiency of SK-FGG was eventually determined using glycerol as the sole carbon source in YNB medium supplemented with 1× 20 mg/L leucine and histidine in a single batch. The conversion experiments were performed under four different conditions of oxygen availability, as mentioned in the Methods section. Significant growth or production of ethanol was not observed under strict anaerobic condition (Table 4). Under micro-aerobic condition, the strain consumed 37.17 g glycerol, with a consumption rate 0.62 g·L·h and production rate 0.25 g·L·h. Acetic acid accumulated at 0.78 g/L under this condition, and the efficiency of ethanol conversion approached 0.42 g^e^/g^g^ (Table 4). Glycerol is consumed more rapidly in a semi-aerobic atmosphere. The consumption rate was > 1 g·L·h, which increased the ethanol production rate to 0.44 g·L·h, resulting in the accumulation of 20.97 g/L ethanol. Acetic acid (2.88 g) accumulated under this condition. As a result, the total conversion reached 0.44 g/g glycerol (Table 4). Under aerobic conditions, glycerol consumption and ethanol production were boosted to 1.29 and 0.5 g·L·h, respectively. The efficiency of ethanol conversion was 0.39 g^e^/g^g^, and the total convertibility was 0.45 g^e^/g^g^, which represents 90% of the theoretical conversion without considering the glycerol utilized in cell formation (Table 4). [“ (Table 4)”]

**Table 4.**
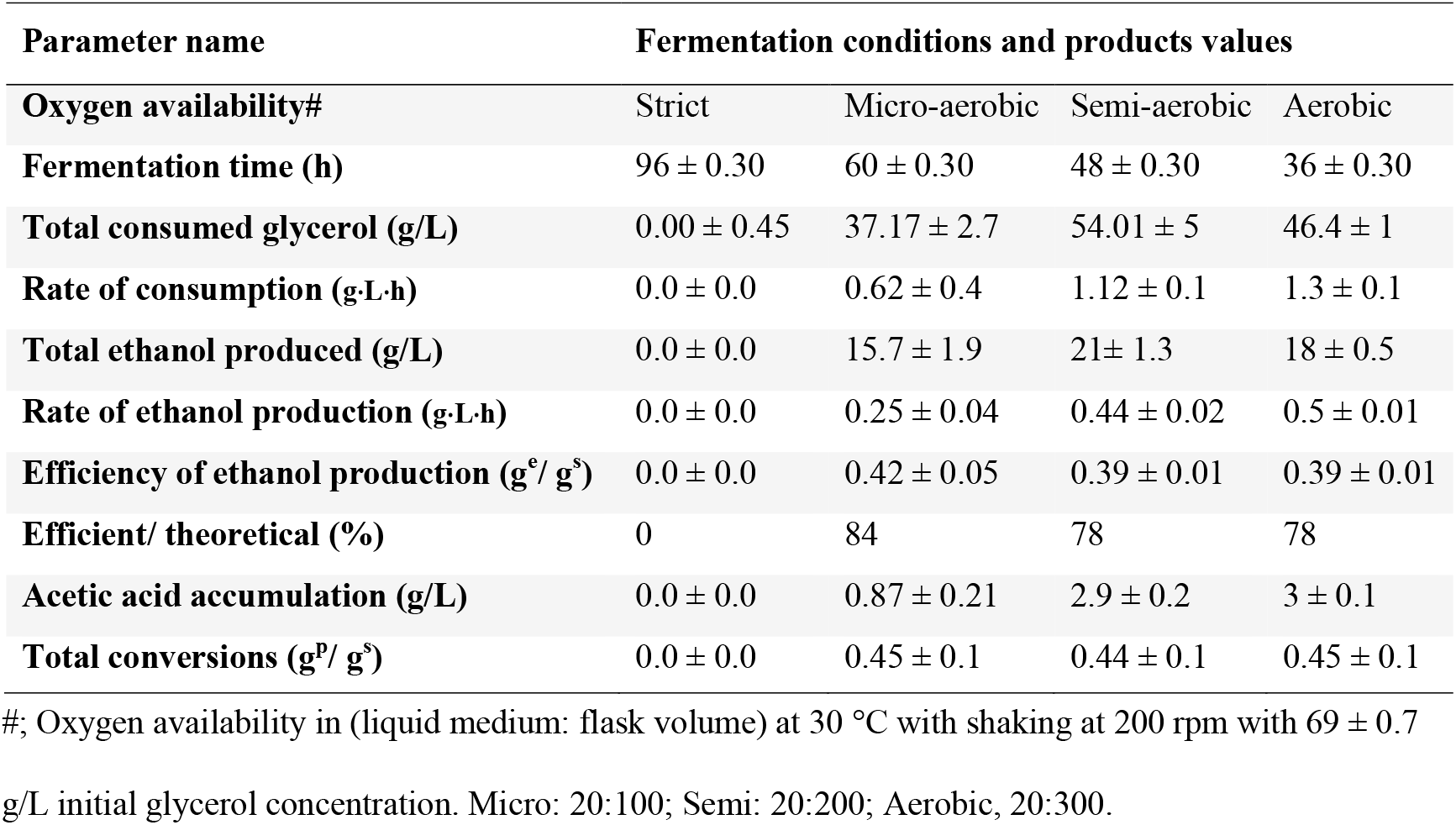
Fermentation characteristics of glycerol as the sole carbon source by strain SK-FGG at different oxygen availabilities.

| Parameter name | Fermentation conditions and products values |  |  |  |
| --- | --- | --- | --- | --- |
| Oxygen availability# | Strict | Micro-aerobic | Semi-aerobic | Aerobic |
| Fermentation time (h) | 96 ± 0.30 | 60 ± 0.30 | 48 ± 0.30 | 36 ± 0.30 |
| Total consumed glycerol (g/L) | 0.00 ± 0.45 | 37.17 ± 2.7 | 54.01 ± 5 | 46.4 ± 1 |
| Rate of consumption (g·L·h) | 0.0 ± 0.0 | 0.62 ± 0.4 | 1.12 ± 0.1 | 1.3 ± 0.1 |
| Total ethanol produced (g/L) | 0.0 ± 0.0 | 15.7 ± 1.9 | 21 ± 1.3 | 18 ± 0.5 |
| Rate of ethanol production (g·L·h) | 0.0 ± 0.0 | 0.25 ± 0.04 | 0.44 ± 0.02 | 0.5 ± 0.01 |
| Efficiency of ethanol production (g<sup>e</sup>/ g<sup>s</sup>) | 0.0 ± 0.0 | 0.42 ± 0.05 | 0.39 ± 0.01 | 0.39 ± 0.01 |
| Efficient/ theoretical (%) | 0 | 84 | 78 | 78 |
| Acetic acid accumulation (g/L) | 0.0 ± 0.0 | 0.87 ± 0.21 | 2.9 ± 0.2 | 3 ± 0.1 |
| Total conversions (g<sup>p</sup>/ g<sup>s</sup>) | 0.0 ± 0.0 | 0.45 ± 0.1 | 0.44 ± 0.1 | 0.45 ± 0.1 |

#; Oxygen availability in (liquid medium: flask volume) at 30 °C with shaking at 200 rpm with 69 ± 0.7

Interestingly, the metabolic engineering here changed the nature of the D452-2 strain from non-grower on glycerol to a grower growing at the rate of 0.045/h (Fig. S2) when starting from an initial inoculum of 0.01, thereby confirming the importance of supplementation of YNB medium with 10× 20 mg/L leucine and histidine in a 50 mL Falcon tube. Growth was also observed during the ethanolic conversion of glycerol before a slight decrease at the onset of the conversion (Fig. S3). A similar tendency was observed during monitoring growth with supplementation of 1× 20 mg/L mg/L leucine and histidine (data not shown). These results suggested that the D452-2 strain was significantly affected by the supplementation levels of histidine and leucine. This observation is supported by a recent study showing a crucial role of amino acid content in cell growth (36). Biosynthesis of NAD^+^ from tryptophan via the kynurenine pathway is well known (37, 38). In addition, nicotinic acid, nicotinamide, quinolinic acid, and nicotinamide riboside can salvage NAD^+^ biosynthesis. In this context, nicotinic acid is auxotrophic under anaerobic conditions in *S. cerevisiae* (38). Together, these limitations may explain the lower production titers, conversion rates, and efficiency in YNB compared to that in the YP medium by the engineered strain. In addition, the other remarkable difference between YNB and YP media is the increased accumulation of acetic acid in YNB (from 14 mg to 53 mg acetic acid/g glycerol) under semi-aerobic conditions (Tables 3, 4). One of the theories assumes a competition between aldehyde dehydrogenase (*ScALD*) and *OpGDH* for NAD^+^; however, further studies are required to clarify this point. The inability to ferment glycerol under strict anaerobic conditions (Table 4) indicates the lack of renovation shuttles in the absence of oxygen and oxidizing agents. Hence, recycling of NADH/NAD^+^ represents the complementary factor for robust oxidation via the DHA pathway and efficient utilization of glycerol to produce bioethanol or other bio-based chemicals. We are currently designing metabolic engineering strategies for glycerol fermentation under anaerobic conditions, while using the high reduction ability of glycerol to improve the fermentation efficiency of lignocellulose sugars. In addition, we intend to determine the amino acids that may play a significant role in the defined media to reach production level of complex medium.

In summary, we presented an efficient model that comprehensively controlled glycerol traffic to ethanol in *S. cerevisiae*. This model of systematic metabolic engineering included integration of the following: (i) robust expression of all genes involved in the DHA pathway; (ii) predominant glycerol oxidation by an oxygen-dependent dynamic of the water-forming NADH oxidase *LlNoxE*, which controls the reaction stoichiometry with the regeneration of the cofactor NAD^+^; (iii) elimination of the first step of both glycerol biosynthesis and glycerol catabolism via G3P. Our study provides an innovative metabolic engineering strategy for re-routing glycerol traffic in *S. cerevisiae*, while tracking ethanol production to levels that have not yet been attained within any other safe model organism, either native or genetically engineered (8, 12, 14, 39-41). The metabolic engineering strategy represents another pivotal step for fermenting glycerol in several promising biorefinery scenarios by co-converting glycerol with the saccharified glucose to ethanol after glycerolysis pretreatment of recalcitrant lignocellulosic biomass.

## MATERIALS AND METHODS

### Strains, primers, cassettes, and plasmid construction

All the strains used in this study (Table 1) were derived from the laboratory strain D452-2 (17), and their original strains are shown in Fig. S1. The plasmids used in this study are listed in Table 5. The primers used are listed in Table S1. The primers used for obtaining the native genes were designed based on the sequences available in the S288C *Saccharomyces* Genome Database. The details of the DNA fragments and cassettes, and the construction of plasmids are mentioned as follows. [“ (Table 5)”]

**Table 5.** Plasmids constructed in this study.

| **Plasmids** | **Relevant genotypes** | **Reference** |
| --- | --- | --- |
| **pPGK-URA3** | *URA3*; *PGK* promoter and terminator | (44) |
| **TDH3-DIT1<sub>d22</sub>-URA3** | *URA3*, *TDH3* promoter and mutated *DIT1<sub>d22</sub>* terminator | This study |
| **TDH3-GDH-DIT1<sub>d22</sub>-URA3** | *URA3*; expresses *OpGDH* | This study |
| **TDH3-NoxE-DIT1<sub>d22</sub>-URA3** | *URA3*; expresses *LINoxE* | This study |
| **pPGK-DAK1-URA3** | *URA3*; expresses *ScDAK1* | This study |
| **pPGK-DAK2-URA3** | *URA3*; expresses *ScDAK2* | This study |
| **pPGK-TPI1-URA3** | *URA3*; expresses *ScTPI1* | This study |
| **pPGK-TPI1-DAK2-URA3** | *URA3*; expresses *ScTPI1*, *ScDAK2* | This study |
| **pPGK-TPI1-DAK2-DAK1-URA3** | *URA3*; expresses *ScTPI1*, *ScDAK2*, *ScDAK1* | This study |
| **pAUR101** | *AUR1-c* | Takara Bio |
| **pAUR101-PGK-RPL41B** | *AUR1-C*, *PGK* promoter-*RPL41B* terminator | This study |
| **pAUR101-FPS1** | *AUR1-C*; expresses *CuFPS1* | This study |
| **pAUR101-FPS1-TPI1-DAK2-DAK1** | *AUR1-C*; expresses *CuFPS1*, *ScTPI1*, *ScDAK2*, *ScDAK1* | This study |
| **pAUR101- M1\*** | *AUR1-C*; expresses *CuFPS1*, *OpGDH*, *ScDAK1*, *ScTPII* | This study |
| **Multiplex pCAS (Addgene #60847)** | Multiplex expresses Cas9 and HDV ribozyme-sgRNA | (42) |
| **Multiplex pCAS/GPD1-1** | Multiplex expresses Cas9 and gRNA to base No. 135 of *ScGPD1* | This study |
| **Multiplex pCAS/GPD1-2** | Multiplex expresses Cas9 and gRNA to base No. 1045 of *ScGPD1* | This study |
| **Multiplex pCAS/GUT1** | Multiplex expresses Cas9 and gRNA to base No. 616 of *ScGUT1* | This study |

\*M1: TEFp-*CuFPS1*-CYC1t; TYS1p-*OpGDH*-ATP15t; TDH3p-*ScDAK1*-d22-DIT1t; FBA1p-*ScTPII*-TDH3t

### Construction of TDH3-DIT1_d22_-URA3, TDH3-GDH-DIT1_d22_-URA3, and TDH3-NoxE-DIT1_d22_-URA3 plasmids

Initially, D452-2 cells picked using a toothpick were resuspended in a polymerase chain reaction (PCR) tube containing 20 µL of 30 mM NaOH (Wako, Osaka, Japan) and heated at 95 °C for 10 min in PCR (Astec thermal cycler-GeneAtlas, Japan). Next, 1 µL of the disrupted cells was used as a template for a 50 µL reaction. High-fidelity polymerase KOD-plus neo (Toyobo, Osaka, Japan) was used to obtain the *TDH3* promoter with an extra region of 52 base pairs (bp) complementary to the upstream sequence of the start codon of *ScGPD1* (fragment 1). The mutated terminator *DIT1*_*d22*_ was purchased from Integrated DNA Technology (IDT; Tokyo, Japan) (21) and then amplified to generate an extra region (46 bp) corresponding to the downstream sequence of the stop codon of *ScGPD1* (fragment 2). Thereafter, the fragments were restricted using XhoI and NotI (Takara, Shiga, Japan) for the first fragment and NotI and SalI for the second fragment. Then, the DNA was purified using the FastGene Gel/PCR extraction kit (Nippon Genetics, Tokyo, Japan). One-step cloning was used to clone the *TDH3* promoter and mutated *DIT1*_*d22*_ terminator into the XhoI/SalI sites of the PGK-URA3 plasmid to construct the TDH3-DIT1_d22_-URA3 plasmid (Table 5). The *OpGDH* gene was purchased from IDT (GenBank accession number XP_018210953.1) (Table S2). NotI/BamHI sites were added to *OpGDH* via PCR and then cleaved by restriction enzymes to form the TDH3-GDH-DIT1_d22_-URA3 plasmid (Table 5). Similarly, *LlNoxE* (accession number AAK04489.1) was prepared as described above. The sequence is also shown in Table S2 and was prepared as described earlier. The plasmid was then cloned into the TDH3*-*DIT1_d22_-URA3 plasmid to assemble the TDH3-NoxE-DIT1_d22_-URA3 plasmid (Table 5).

### Construction of the multiplex pCAS-gRNA-CRISPR system

The original multiplex pCAS-gRNA system was a gift from Prof. Jamie Cate (42). The online tool https://crispr.dbcls.jp/ was used for the rational design of the CRISPR/Cas target (43). The efficiency of the target design was also confirmed using https://chopchop.cbu.uib.no/. Accordingly, a highly specific guide, 20 base pairs (bp) before the protospacer adjacent motif (PAM), was selected and used to design the primers (Table S1). Instead of a single-step amplification of the entire multiplex plasmid, which could result in mutations, two rounds of polymerization were used to produce only universal scaffolds. In the first round, the two fragments were synthesized separately. The first fragment encompassed the upstream region of the guide RNA (gRNA) scaffold, while the second one encompassed the downstream region (Fig. S4). After purifying each DNA segment, the overlapping gRNA (20-nucleotide guide sequence) was used to generate overhangs during the second round of PCR (Fig. S4). During the second PCR, KOD-plus neo, pCAS F, and R primers were used with 6 picograms of each DNA segment as a template. Then, the unified DNA fragment was cleaved by SmaI/PstI to form SmaI-universal scaffold-PstI. Subsequently, the universal scaffold was cloned into a multiplex plasmid, and its analogous part was detached. As a result, a new multiplex pCAS-gRNA plasmid was formed. These steps were repeated for the construction of multiplex pCAS-gRNA plasmids targeting *ScGPD1* and *ScGUT1* (Table 5). The sequences of the new SmaI-universal scaffold-PstI in the constructed plasmids were confirmed via sequencing.

### Construction of pPGK-DAK1-URA3, pPGK-DAK2-URA3, pPGK-TPI1-URA3, pPGK-TPI1-DAK2-URA3, and pPGK-TPI1-DAK2-DAK1-URA3 plasmids

*DAK1, DAK2*, and *TPI1* were obtained from genomic DNA of the parent strain D452-2, as mentioned above. The Xhol site of *ScDAK2* was disrupted by incorporating a silent mutation before cloning. The ends of the genes were cleaved using the restriction enzymes, sites of which were present in the supplemented primers (Table S1, section 1). First, each gene was separately cloned in the pPGK-URA3 plasmid (44) under the *PGK* promoter and terminator (Table 5). After cloning, the sequence of the genes was confirmed using the primers listed in Table S1. To construct the pPGK-TPI1-DAK2-URA3 plasmid, the XhoI/SalI-*ScDAK2* cassette was inserted in the SalI site of the pPGK-TPI1-URA3 plasmid after dephosphorylation of the site. Since the previous ligation of SalI with XhoI altered the sequence of the ligation site, preventing its cleavage and detachment of the cloned cassettes, XhoI/SalI-*ScTPI1* or *ScDAK1*, we reused the SalI site during the cloning of the XhoI/SalI-*ScDAK2* cassette. The plasmid pPGK-TPI1-DAK2-DAK1-URA3 was then constructed (Table 5).

### Construction of pAUR101-PGK-RPL41B, pAUR101-FPS1, and pAUR101-FPS1-TPI1-DAK2-DAK1 plasmids

*Candida utilis* (NBRC 0988) was obtained from the National Biological Resource Center (NBRC) of the National Institute of Technology and Evaluation (Japan) and was used as a source of the glycerol facilitator (*CuFPS1*). The sequence of *CuFPS1*, which was included in the deposited gene bank under the accession number BAEL01000108.1, is shown in Table S2, along with its expression system, including the PGK promoter and RPL41B terminator. To construct the pAUR101-PGK-RPL41B plasmid, the pAUR101 vector (Takara, Shiga, Japan) was subjected to one step of cloning of the SmaI-PGK-Not1 and NotI-RPL41B-SacI fragments into its SmaI-SacI sites. Thereafter, *CuFPS1* was cloned into a dephosphorylated NotI site of pAUR101-PGK-RPL41B to assemble the pAUR101-FPS1 vector. The direction of cloning was confirmed using PCR. Then, a detached set of cassettes, *ScTPI1, ScDAK2*, and *ScDAK1*, obtained via Xhol-SalI digestion from the previously constructed plasmid, pPGK-TPI1-DAK2-DAK1-URA3, was cloned into the dephosphorylated SalI site of the pAUR101-FPS1 vector to assemble the pAUR101-FPS1-TPI1-DAK2-DAK1 plasmid (Table 5).

### Construction of the M1 and pAUR101-M1 plasmids

All fragments constituting this module were first obtained separately via PCR (Fig. 3). *CuFPS1, OpGDH*, and the mutated *DIT1*_*d22*_ terminator were amplified from their synthetic DNA stocks, while the other fragments were amplified from the genomic DNA of the D452-2 strain. Following the generation of 12 fragments via PCR, the products were electrophoresed on 1% or 2% agarose gel, excised according to the size of fragments, and purified using an extraction kit (Nippon Genetics, Tokyo, Japan). The highly purified fragments were used for assembly using the Gibson assembly master mix (New England Biolabs, Tokyo, Japan). First, three consecutive segments were joined seamlessly according to the manufacturer’s protocol (Gibson), followed immediately by PCR for the next consecutive three segments, which were then purified again from agarose gel. Six sequential segments were assembled via Gibson assembly and PCR to assemble the whole module M1: TEFp-*CuFPS1*-CYC1t; TYS1p-*OpGDH*-ATP15t; TDH3p-*ScDAK1*-DIT1_d22_t; FBA1p-*ScTPI1*-TDH3t (Fig. 3). The SacI site was added upstream of module M1, while the SmaI site was added downstream for cloning into the pAUR101 vector to form pAUR 101-M1 (Table 5). Finally, the vector pAUR 101-M1 was transformed into *Escherichia coli* as described above, and the accurate structure of M1 was confirmed by sequencing the whole module M1 in the pAUR-M1 vector.

### Transformation and recombination of strains

All constructed plasmids were transformed in *E. coli* NEB 10-beta via the heat shock method of transformation according to the manufacturer’s instructions (New England Biolabs, Tokyo, Japan). All plasmids were extracted using the QIAprep Spin miniprep kit (QIAGEN, Hilden, Germany). All DNA measurements were performed using a BioSpec-nano (Shimadzu, Japan). Plasmids and DNA fragments were stored at 20 °C. Yeast transformation was performed using the Fast™ yeast transformation kit (G-Biosciences, USA) to integrate the linear pAUR101 vector. BsiWI (New England Biolabs, Tokyo, Japan) was used to linearize the *AUR1-C* gene before the transformation, which was integrated in the *AUR1* locus of *S. cerevisiae* via homologous recombination. Positive colonies were selected on aureobasidin A (Takara, Shiga, Japan), and further confirmed via PCR. The same method and kit were used to transform the linearized pPGK-URA3 plasmid, in which *URA3* was linearized with NcoI (Takara, Shiga, Japan). The transformants were grown on Yeast Nitrogen Base (YNB) medium supplemented with 20 mg/L leucine and histidine; the fast-growing colonies were confirmed using PCR and then screened for glycerol consumption and ethanol production. *ScGPD1* with *OpGDH* were replaced via homologous repair of the double-strand break, which was induced by CRISPR Cas 9 (45). Similarly, *ScGUT1* was substituted by the M1 module. The homologous regions were further extended via PCR during double-strand DNA repair. Positive colonies were confirmed via PCR using primers located on the inserted cassettes and regions upstream and downstream of the flanking recombined loci. The recombinant cells were re-confirmed after loss of the pCAS multiplex plasmids. All the recombinant strains and their genotypes are listed in Table 1.

### Cultivation of yeast cells

Yeast peptone dextrose (YPD) medium was routinely used for the maintenance and cultivation of strains listed in Table 5 on solid agar or in liquid media. All liquid cultivation was conducted under micro-aerobic conditions (10 mL culture in 50 mL Falcon tubes with 200 rpm of orbital shaking (TAITEC, BioShaker BR-42FL, Japan) and approximately 45° sitting angle) overnight at 30 °C. The screw caps of the tubes were closed at a level that permitted CO_2_ escape. Under these conditions, the optical densities (OD_600_) of the strains varied between 6 and 9 (Fig. S5). The cell densities were monitored at 600 nm after ten-fold dilution using a spectrophotometer (AS ONE, Japan). YPD_15_G_70_ medium [(w/v) 1.5% glucose and 6% glycerol] was used for preculturing the cells, as shown in Tables 2 and 3. In addition, it was used during cultivation of the cells, which were used for preparing the cell-free extracts and determining the NAD^+^/NADH ratios. YNB medium supplemented with 1× 20 g/L glucose, 20 mg/L leucine, and histidine was used to culture the recombinant strains harboring active *URA3* during or after the transformation.

### Preparation of cell-free extract

Proteins were extracted as described previously (22, 46, 47) with some modifications as follows: 10 mL cell culture of approximately OD_600_ = 5 was harvested via centrifugation at 700 × *g* for 2 min at 4 °C. The cell pellets were washed twice with 20 and 1 mL of 100 mM HEPES buffer (pH 7.4). Then, the cell pellets were lysed in 1 mL HEPES buffer supplemented with 1 mM MgCl_2_ and 10 mM 2-mercaptoethanol with approximately 400 mg glass beads in a 2 mL Eppendorf tube. The lysis was accomplished by vigorous shaking using a bench vortex with six time intervals on ice for 30 s. The crude proteins were separated from the glass beads and cell debris via two rounds of centrifugation at 22,300 × *g* at 4 °C for 5 min. The total protein concentration of the supernatant was estimated using Quick Start™ Bradford (Bio-Rad, USA) and bovine serum albumin standards (Novagen, USA) at 595 nm using Infinite M200 (Tecan, Switzerland).

### Enzyme assays

The specific activity of glycerol dehydrogenase was assayed by monitoring the changes in NADH absorbance at 340 nm (UV-2700; UV-VIS spectrophotometer, Shimadzu, Japan). One milliliter mixture of 79 mM HEPES buffer (pH 7.4), 10 µL crude extract, and 10 mM NAD^+^ were mixed in the quartz cuvette and incubated for 30 s to complete any side reaction and obtain a stable baseline. The reaction was initiated by adding 100 µL of 1 M glycerol and incubating for 1 min. The activity of *ScTPI1* was assayed as described previously (48), with some modifications. The reaction mixture consisted of 100 mM (pH 7.53) triethanolamine hydrochloride (Sigma-Aldrich), 10 mM dihydroxyacetone phosphate (DHAP) hemimagnesium salt (Sigma-Aldrich), 100 µL crude extract, and 5 mM NAD^+^ as a starting point for the reaction, which lasted 400 s. The main changes in absorbance were detected after approximately 100 s. Changes in the absorbance from 200 s to 260 s were used to calculate the activities. Dihydroxyacetone kinase was assayed using a universal kinase activity kit according to the manufacturer’s instructions (R&D systems, EA004, USA).

### Determination of the intracellular concentration of NADH/NAD^+^

NADH/NAD^+^ was estimated as reported previously (24). Twenty microliters of cultivated cells of OD_600_ = 5 were harvested via centrifugation at 1600 × *g* for 2 min at 4 °C. The NADH/NAD^+^ ratio was determined per the instructions of the EnzyChrom™ NAD^+^/NADH assay kit (BioAssay Systems (E2ND-100)).

### Fermentation

Fermentations were performed with orbital shaking (TAITEC, BioShaker BR-42FL, Japan) at different levels of oxygen availability in the flask based on the ratio of liquid medium (mL): Erlenmeyer flask volume (mL), namely, micro-aerobic (20:100), semi-aerobic (20:200), and aerobic (20:300). The cell pellets were obtained from approximately 20 mL cell culture (OD_600_ = 4), which was harvested from appropriate volumes of preculture via centrifugation at 1,600 × *g* for 5 min and then washed with 20 mL Milli-Q water. The alcoholic conversion of glycerol was conducted at 30 °C with agitation speeds of 180 or 200 rpm. The YP and YNB media were supplemented with glycerol or glucose of different initial concentrations, which listed in Fig. 2, Table 3, and the legend of Table 4, in accordance with those obtained after the glycerolysis of biomass (16). The initial concentrations of glycerol or glucose are shown in Tables 3 and 4 and Fig. 2. For fermentation under strict anaerobic conditions, 50 mL Mighty vials were sterilized with a precision seal septum cap prior to inoculation of the cell pellets and YNB medium. Then, nitrogen gas was flushed into the medium through Terumo needles, which were also used for sampling.

### Fermentation analysis

The samples (100 µL) were picked from fermentative flasks under sterile conditions, diluted with 900 µL Milli-Q water in Eppendorf tubes, and mixed before centrifugation at 22,300 × *g* for 5 min. Subsequently, the supernatants were decanted using a 1 mL syringe and filtered through a 0.45 µm hydrophilic filter (PTFE) directly into new 2 mL high performance liquid chromatography (HPLC) glass vials. Analyses were performed using HPLC (Shimadzu, Japan) on an Aminex HPX-87H column (Bio-Rad Laboratories, Hercules, CA, USA) connected to refractive index (RID-10A; Shimadzu) and prominence diode array (SPD-M20A; Shimadzu) detectors using 5 mM H_2_SO_4_ as the mobile phase at 50 °C with a flow rate of 0.6 mL/min. Glucose, glycerol, ethanol, acetic acid, pyruvate, succinic acid, and acetaldehyde were monitored using RID. DHAP and DHA levels were quantified using a prominence diode array detector (SPD-M20A).

## Data availability

All necessary data required to assess our findings are available in this manuscript or its supplementary data. Further details related to this article may be requested from the authors.

## Author contributions

S.M.R.K. conceived the research idea, and planned the experiments. S.M.R.K. also provided information regarding procurement of strains, chemicals, and toolboxes for genetic engineering, performed the experiments, and analyzed the results. S.M.R.K. wrote, revised, and submitted the manuscript. T.W. was responsible for all financial support and provided all the chemicals and equipment. T.W. was involved in formulating the research idea, planning and organization, discussion of the results, and manuscript revision and submission.

